# Comparative pangenomics unveils distinct host adaptation levels and conserved biosynthetic potential in microbiome *Clostridia*

**DOI:** 10.1101/2025.02.12.637840

**Authors:** Lucas De Vrieze, Jan Aerts, Joleen Masschelein

## Abstract

To thrive in diverse ecological niches, bacteria adopt various lifestyles, that range from living freely in the soil to forming close associations with human and animal hosts. However, the impact of these adaptation processes on their genomes and metabolisms remains largely unexplored beyond the genus level. Investigating these evolutionary dynamics at higher taxonomic levels can enhance our understanding of the relationship between host adaptation and their functional capabilities. Here, we examine the evolutionary trajectories and metabolic capabilities of the *Clostridia* class, which displays a variety of lifestyles and is of high importance for industry, medicine and microbiome research. First, we uncover that the clostridial orders have significantly different adaptation rates. Second, we show that the *Oscillospirales* order has undergone extensive genomic and functional specialisation toward a host-associated lifestyle, while the *Lachnospirales* order tends to be at a lower level of host association, retaining a remarkably high number of free-living trait genes and a high degree of metabolic versatility. Third, we reveal substantial differences in genomic architecture and metabolic versatility between the clostridial orders and link these to the progressing stages of host adaptation. Additionally, we identify widely conserved biosynthetic gene clusters, highlighting untapped biosynthetic potential of evolutionary significance. Hence, the beyond-genus level analyses in this study provide valuable new insights into bacterial adaptation with broad implications for evolutionary biology, microbiome research and biotechnology.

## Introduction

Microorganisms continuously interact with their environment, adapting not only through short-term regulatory and metabolic responses, but also through long-term genomic changes. The continuous acquisition, mutation and deletion of genes is closely tied to environmental selection pressures, giving rise to various lifestyles that vary in their dependence on external factors. These selection pressures drive evolutionary processes, such as host adaptation and genome streamlining, which can comprehensively reshape microbial genomes. This, in turn, has a profound impact on their functional and metabolic versatility, and hence their ability to survive in different ecological niches (Dufresne, Garczarek, and Partensky 2005; Siguier, Gourbeyre, and Chandler 2014; Marais, Calteau, and Tenaillon 2008).

Species belonging to the *Clostridia* class exemplify this evolutionary and ecological diversity. They exhibit a remarkable variety of lifestyles and phenotypes, ranging from thermophiles and psychrophiles to acidophiles and pathogens. They occupy diverse environments, including soil, sediments and the intestinal tracts of humans and animals. This versatility underlies their increasing industrial and medical relevance, exemplified by industrially important species like *C. acetobutylicum* and *C. beijerinckii* (Birgen et al. 2021; Patakova et al. 2019; Roberts et al. 2010) and notorious pathogens like *C. botulinum, C. tetani* and *C. difficile* (Cohen et al. 2017; Smith, Hill, and Raphael 2015; Sandhu and McBride 2018). Moreover, *Clostridium* species were the first anaerobes shown to produce biosynthetic gene cluster (BGC)-encoded secondary metabolites with potent biological activities, reflecting their relevance for natural product discovery (Pahalagedara et al. 2020; Letzel, Pidot, and Hertweck 2013; Behnken and Hertweck 2012; Sugimoto et al. 2019).

Despite the abundance, diversity and relevance of *Clostridia*, only few comparative studies have explored their evolutionary history and ecological adaptations beyond the species level. Previous higher-level comparative studies focused on a few individual species or a single genus at most. For example, a study of *Clostridium sensu stricto* found evidence of a highly plastic genome, harbouring a relatively large metabolic core and many specialised genetic capabilities that underpin the mainly free-living nature of this genus (Udaondo, Duque, and Ramos 2017). Another study mainly focusing on *Clostridium sensu stricto* identified divergent evolution of the shikimate pathway in pathogenic clostridia and extensive mobility among their toxin genes (Cruz-Morales et al. 2019). Lastly, a study of the gut-associated genus *Blautia* revealed that many gene exchange events involve metabolic functions, likely reflecting adaptation to the dynamic host environment, and suggested that such exchanges may have caused a metabolic dependence on other gut bacteria (Maturana and Cárdenas 2021). While informative, these studies do not provide broader class-level insights that allow cross-genus comparison.

In this work, we present a comprehensive study of the interplay between host adaptation, genome evolution and metabolic capacity for the taxonomic class of *Clostridia*. By combining pangenomic and phylogenetic analyses, we characterise the evolutionary dynamics within different orders and infer the lifestyles of their ancestral lineages. In parallel, we assess the metabolic versatility of each species and link this to their evolutionary trajectory. We also screened all genomes for genomic hallmarks and mobile genetic elements (MGEs) associated with progressing host adaptation. Finally, we zoom in on secondary metabolite BGCs, assessing their conservation and evolutionary dynamics across the class, and reveal biosynthetic features of potential ecological and biotechnological significance.

## Materials and methods

### Data gathering, QC and filtering

All genome assemblies were retrieved from RefSeq using the search query “Clostridia” in the NCBI Datasets CLI (v16.2.0) (O’Leary et al. 2024), excluding MAGs and atypical assemblies (# = 10599 as of 22/05/2025). The resulting set was filtered by number of contigs (<= 200), N50 (>= 50 kb), completeness (>= 90%) and contamination (<= 10%), and dereplicated at species level using skDER (v1.3.1) (Salamzade, Kottapalli, and Kalan 2025) (ANI cutoff: 96%, AF cutoff: > 50%; **Add. File 2: Figure S1**), resulting in 1010 representative assemblies. For consistency, all assemblies were reannotated with Bakta (v1.9.1) (Schwengers et al. 2021) and taxonomically verified using GTDB-tk (v2.4.1) (Chaumeil et al. 2022). To ensure sufficient statistical power in downstream analyses, only taxonomic orders with at least 30 representative species were retained.

### Metabolic versatility analysis

The metabolic versatility of all taxa was assessed by KEGG module completeness as determined by the MicrobeAnnotator (v2.0.5) pipeline (Ruiz-Perez, Conrad, and Konstantinidis 2021) in light mode at default settings. The module completenesses were clustered using Ward’s hierarchical clustering and plotted using in-house Python scripts. Optimal leaf ordering was enabled while plotting the clustermap.

### Pangenome construction and analysis

A pangenome was constructed for the full genome set using PPanGGOLiN (v2.0.5) (Gautreau et al. 2020) with gene defragmentation enabled. To account for the high diversity in this class-level genome set, clustering was carried out with a relaxed sequence identity threshold of 40% and an alignment coverage cutoff of 40%, while the pangenome was partitioned using a core genome threshold of 90% and an accessory genome threshold of 0.099% (or 1/1010), as appeared most appropriate from our parameter sweep (**Add. File 3**). Presence/absence matrices for the different clostridial orders were extracted from the full class-level matrix by selecting for gene families present in at least one taxon from that order. The same pangenome partitioning thresholds were applied for consistency.

### Phylogeny construction

A phylogenetic tree for the full clostridial genome set was constructed with PhyloPhlAn (v3.0.67) (Asnicar et al. 2020) with a custom gene marker database derived from the representative protein sequences of the earlier identified core gene families. The tree was rooted using *B. subtilis* 168 as outgroup (RefSeq: GCF_000009045.1). The phylogeny was computed at medium diversity setting using DIAMOND (Buchfink, Reuter, and Drost 2021) as the mapping algorithm, MAFFT (Katoh and Standley 2013) as the alignment algorithm, Trimal (Capella-Gutiérrez, Silla-Martínez, and Gabaldón 2009) as the alignment trimmer, and IQ-TREE2 (Minh et al. 2020) as the tree constructor. 10.000 ultrafast bootstraps (Minh, Nguyen, and von Haeseler 2013) were requested by manually adding the “-bb 10000” flag to the IQ-TREE entry in the PhyloPhlAn configuration file. The outgroup was trimmed from the rooted phylogeny using the R package *ape*, after which the tree was visualised using iTol (v6.9) (Letunic and Bork 2024) and ggtree (v3.11.1) (Yu et al. 2017).

### Functional annotation of the pangenome

The protein family reference sequences of the pangenome were functionally annotated using eggNOG-mapper (v2.1.12) (Cantalapiedra et al. 2021). COG category counts and fractions were calculated using pandas (v2.2.1) and plotted using ggplot2 (v3.5.1).

### Genomic divergence rate analysis

A genomic divergence rate analysis estimates the rate of gene exchange within a clade and statistically compares these rates for significant differences. It requires the phylogenies of each examined clade to have an identical time scale. Therefore, a subtree was extracted for each order from the full phylogeny using *ape*. Gene exchange (GE) models were inferred for each order using *Panstripe* (Tonkin-Hill et al. 2023). All options were left at their defaults, except that Gaussian distributions were employed to fit the underlying generalised linear models (GLMs). Cumulative GE plots and statistical comparisons of the GE models were carried out using the appropriate *Panstripe* functions.

### Ancestral state reconstruction

To track the evolutionary origin of gene exchanges, the ancestral states of all gene families were reconstructed from the full core genome phylogeny and the full pangenome presence/absence matrix. This was done according to Wagner parsimony with equal penalty scores for both gene gains and losses using Count (v10.4) (Csurös 2010). KOG annotations of the genes exchanged at the tree nodes of interest were retrieved from the earlier made eggNOG annotations and visualised using in-house R scripts.

### Mobilome analysis

Pseudogenes were detected using Pseudofinder (v1.1.0) (Syberg-Olsen et al. 2022) at default settings. The NCBI Protein database filtered for taxon ID 186801 (the *Clostridia* class) was used as reference database and indexed using DIAMOND (v2.0.14). Insertion elements (ISEs) were inferred using ISEscan (v1.7.2.3) (Xie and Tang 2017) at default settings. Biosynthetic gene clusters (BGCs) were identified using antiSMASH (v8.0.0) (Blin et al. 2025) at default settings. Prophages were found by running PHASTEST (Wishart et al. 2023) in lite mode using the image from Docker Hub (digest 2477a2f3b6b8). Prophage host graphs were made using *supervenn* (v0.5.0). All mobile element counts were normalised by dividing by the genome length and statistically compared using the Brunner-Munzel test (α = 0.05) in scipy (v1.15.3). Significance annotations were added using *statannotations* (v0.6).

### BGC clustering analysis

All BGCs identified during the mobilome analysis were clustered using BiG-SCAPE (v1.1.8) (Navarro-Muñoz et al. 2020), adding MIBiG reference BGCs (flag ‘--mibig’), mixing all BGC classes (flag ‘--mix’), and including singletons (flag ‘--include_singletons’). A BiG-SCAPE distance cutoff value of 0.3 was applied as appeared most appropriate from a parameter sweep (**Add. File 4**) and the resulting similarity networks. To integrate phylogroup metadata in CytoScape (v3.10.2), the unidirectional network links in the output network files of BiG-SCAPE were annotated by order or MIBiG and converted into bidirectional links by adding the reverse link. BGC gene alignments were generated using clinker (v0.0.28) (Gilchrist and Chooi 2021).

## Results

### Genome dataset composition

To examine how host adaptation has shaped genome architecture, metabolic potential and biosynthetic capacity in the *Clostridia* class, we first assembled a comprehensive dataset by downloading all available *Clostridia* genomes from NCBI. Following quality control and species-level dereplication, the final dataset comprised 1010 genomes, sampled from various isolation sources, including open ecosystems as well as host-associated niches, such as human and animal microbiomes. After taxonomic classification using GTDB-tk, we selected four orders that together cover a large fraction of the *Clostridia* class while being sufficiently large for strong statistical power (# >= 30) (**Add. File 2: Figure S2**): *Clostridiales, Lachnospirales, Oscillospirales* and *Peptostreptococcales*. The *Clostridiales* mainly comprise free-living species belonging to *Clostridium sensu stricto*. The *Lachnospirales* and the *Oscillospirales* contain prevalent members of the human gut microbiome (the historical *Clostridium XIVa* and *Clostridium IV* species groups, resp.), while the *Peptostreptococcales* are also commonly found as part of various microflora in the human body (e.g. *C. difficile*).

### Pangenome partition sizes and functional annotation indicate high genome plasticity

To investigate evolutionary dynamics, we first captured the gene conservation in all genomes by constructing a pangenome for the full genome dataset. Using a set of clustering and partitioning thresholds carefully selected after a parameter sweep (**Add. File 3**), the pangenomes for each assessed group of taxa were built and partitioned (**Table 1**). The recurringly large shell and cloud partitions suggest that there is a high level of diversity among all orders, meaning that each genome is likely to harbour unique gene families. This finding highlights the high level of genomic plasticity that is typical for species living in highly dynamic environments like open ecosystems or human-associated niches.

**Table 1.** Number of gene families in the pangenome partitions for each examined order, both in absolute numbers and relative to the total number of gene families in each order’s pangenome.

|  | **All genomes**<br>**(# = 1010)** | ***Clostridiales***<br>**(# = 209)** | ***Lachnospirales***<br>**(# = 349)** | ***Oscillospirales***<br>**(# = 164)** | ***Peptostreptococcales***<br>**(# = 94)** |
| --- | --- | --- | --- | --- | --- |
| **Core (> 90%)** | 353<br>0.09 % | 690<br>0.79 % | 574<br>0.36 % | 465<br>0.59 % | 444<br>0.81 % |
| **Shell**<br>**(0.1 % - 90%)** | 158,627<br>39.78 % | 39,179<br>44.64 % | 63,580<br>40.06 % | 30,325<br>38.43 % | 21,451<br>39.04 % |
| **Cloud (< 0.1%)** | 239,797<br>60.16 % | 47,903<br>54.58 % | 94,546<br>59.58 % | 48,109<br>60.98 % | 33,053<br>60.15 % |

After establishing the pangenome partitions, we examined the functional differences between the orders (**Figure 1**). The functional profile is similar for most core genomes and is dominated by ribosomal proteins (COG category J). A notable difference is the higher proportion of signal transduction genes in the core genome of *Clostridiales* (COG category T), consistent with a free-living lifestyle requiring more complex sensing of the environment and more specialised responses. In almost all orders, the majority of the shell and cloud gene families have no known function (COG categories S and -), suggesting the presence of mobile genetic elements (MGEs), highly specialised functions, and/or rapidly evolving genes that fall outside the scope of our current understanding.

**Figure 1.**
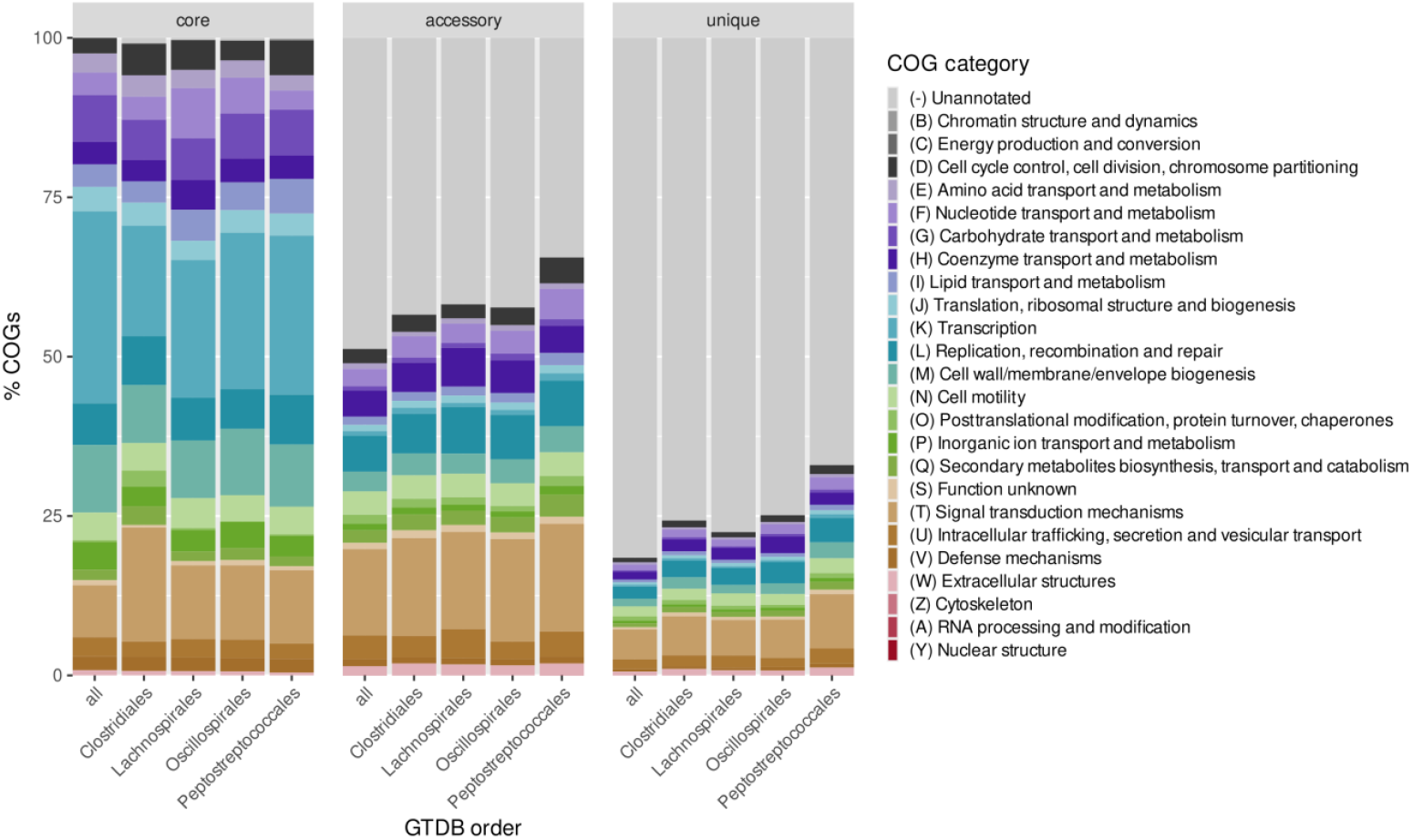
COG functional annotation profiles for all pangenome partitions of each examined clostridial order. Fractions are referenced to the number of gene families in a partition. The functional profiles are similar, except for a larger share of signal transduction genes in the core genome of the *Clostridiales* order, which indicates a free-living lifestyle. For the exact category fractions and counts, see Add. File 8.

### KEGG pathway completeness analysis uncovers two distinct levels of metabolic versatility

A KEGG pathway completeness analysis gauges the metabolic versatility of an organism by assessing the completeness of metabolic pathway modules (KEGG Pathway modules) based on the genomic presence of their constituent genes. Remarkably, two distinct levels of pathway completeness were observed for the full genome dataset. 394 out of 1010 organisms exhibit substantially higher pathway completeness, indicative of more versatile metabolic capabilities (**Figure 2**). This difference is not driven by a limited number of pathways, but is the result of a genome-wide trend, as multiple modules tend to be more complete. This aligns with earlier observations of a remarkably high metabolic versatility within the *Bacillota* phylum, especially within the *Clostridium* genus (Pascal Andreu et al. 2023; Vieira-Silva et al. 2016).

**Figure 2.**
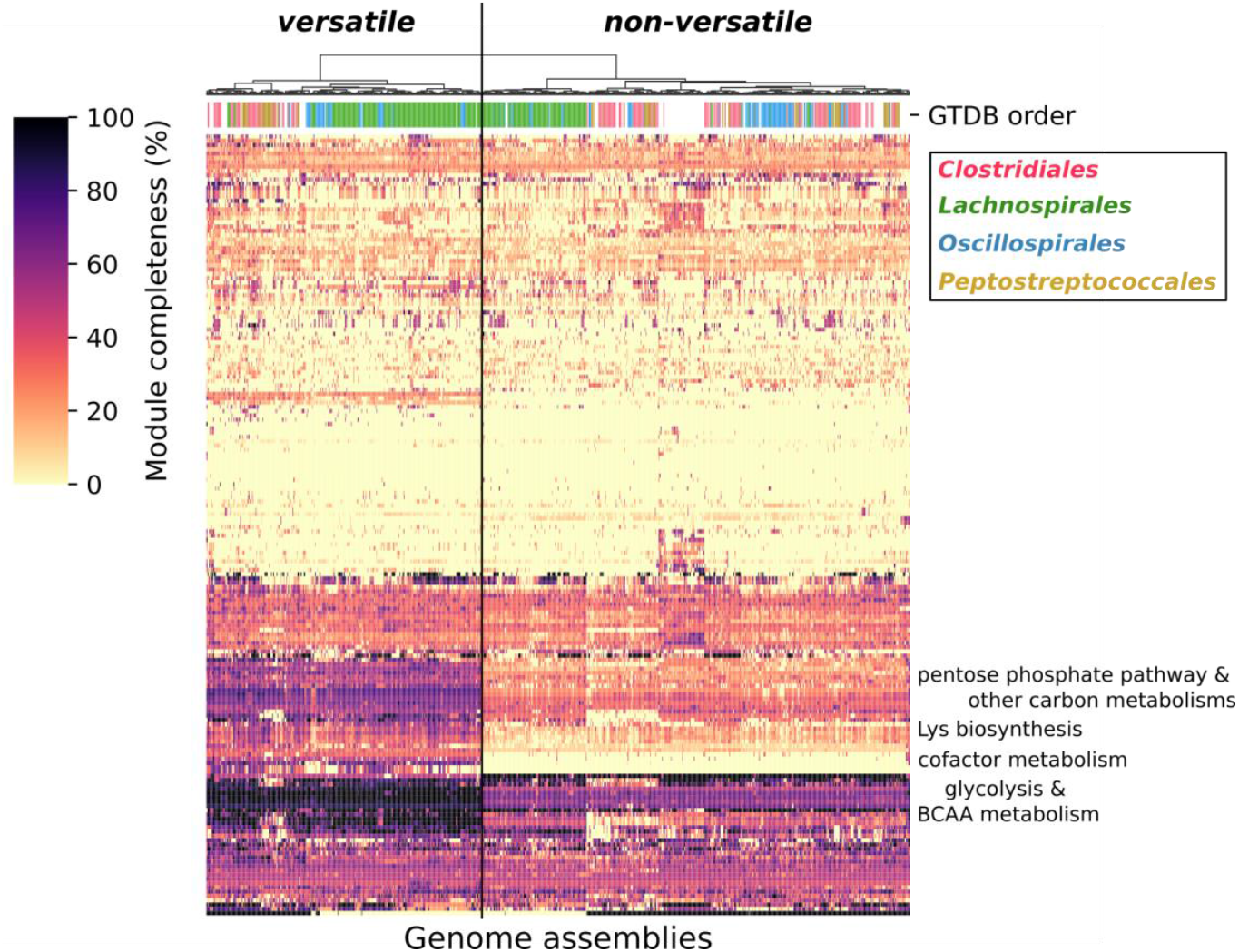
Ward’s clustered heatmap of KEGG module completeness, annotated by taxonomic order. Metabolically versatile genomes recurringly show a general trend in higher module completenesses. In summary, the key differing modules comprise glycolysis-related and BCAA biosynthesis pathways, as well as Lys biosynthesis, the pentose phosphate pathway and other carbon metabolisms. For a figure with all KEGG module labels, see Add. File 2: Fig. S5.

Interestingly, the *Lachnospirales* order has a higher fraction of versatile species than the other orders (55% vs. 25% - 35%; **Add. File 2: Supp. Fig. S3**). Moreover, it is the only order comprising genera and lineages that are (almost) entirely metabolically versatile (*Enterocloster, Lacrimispora, Butyvibrio*, one of two *Blautia* lineages) This metabolic versatility appears independent of genome size, except for the *Clostridiales* order (**Add. File 2: Supp. Fig. S4**). Multiple clostridial genera contain both versatile and non-versatile species, confirming an intra-genus heterogeneity that has previously been reported (Vieira-Silva et al. 2016).

The most pronounced differences in module completeness are found in core metabolic processes, including central carbon metabolism (glycolysis, pentose phosphate pathway) and branched-chain amino acid and Lys biosynthesis. Metabolically versatile *Lachnospirales* species thus have a higher degree of metabolic independence than typically expected for host-associated bacteria. This more developed primary metabolism may, in turn, support a more extensive secondary metabolism, providing additional adaptive advantages.

### Genomic divergence rate analysis reveals significantly different evolutionary dynamics

The higher metabolic versatility of the *Lachnospirales* suggests a distinct evolutionary trajectory compared to the host-associated *Oscillospirales* and *Peptostreptococcales* orders. To assess whether this difference indeed reflects distinct evolutionary dynamics, gene exchange (GE) models were fitted for each order separately and statistically compared to each other using *Panstripe*.

*Panstripe* uses a compound generalised linear model (GLM) to relate phylogenetic branch lengths to the number of genes exchanged at those branches. Using an additional GLM term, internal nodes of the tree can be differentiated from terminal ones, yielding core and tip gene exchange rates, respectively. Exchange rates that are significantly different from zero, signify an association between branch length and the number of gene exchange events, and thus active gene exchange. The core rate represents the long-term gene exchange rate shaping the overall phylogeny structure, while the tip rate captures short-term genomic dynamics, such as transposon- or phage-mediated gene mobility. However, in highly diverse genome sets, such as the dataset analysed here, orthologous gene families may have been split into multiple families during pangenome construction, artificially inflating both core and tip gene exchange rates. Consequently, *Panstripe* models here a more general genomic divergence (GD) rate, that captures both intrinsic gene evolution as well as gene exchanges, rather than only the latter. Nevertheless, we argue that this more general genomic divergence rate is still well-suited to address our research question, as it provides a measure of the pace at which genomes have been evolving.

GD modelling revealed several interesting differences in evolutionary dynamics between the four orders (**Figure 3**). First, the *Oscillospirales* have the lowest core divergence rate, which is consistent with the observation that streamlined genomes of host-associated species evolve at a reduced rate (Dufresne, Garczarek, and Partensky 2005; Marais, Calteau, and Tenaillon 2008). In contrast, the *Lachnospirales* have significantly higher core and tip divergence rates, indicating a higher level of both long-term gene divergence and short-term gene mobility, respectively. This pattern is unexpected for a host-associated order and suggests that *Lachnospirales* are still undergoing active host adaptation (see also **Figure 5G**) (Siguier, Gourbeyre, and Chandler 2014). The tip divergence rate of the *Peptostreptococcales* is higher as well, but the core rates are not. This may reflect increased gene mobility at the onset of host adaptation, or, alternatively, a late stage of host conversion, in which long-term divergence has slowed and residual gene mobility persists.

**Figure 3:**
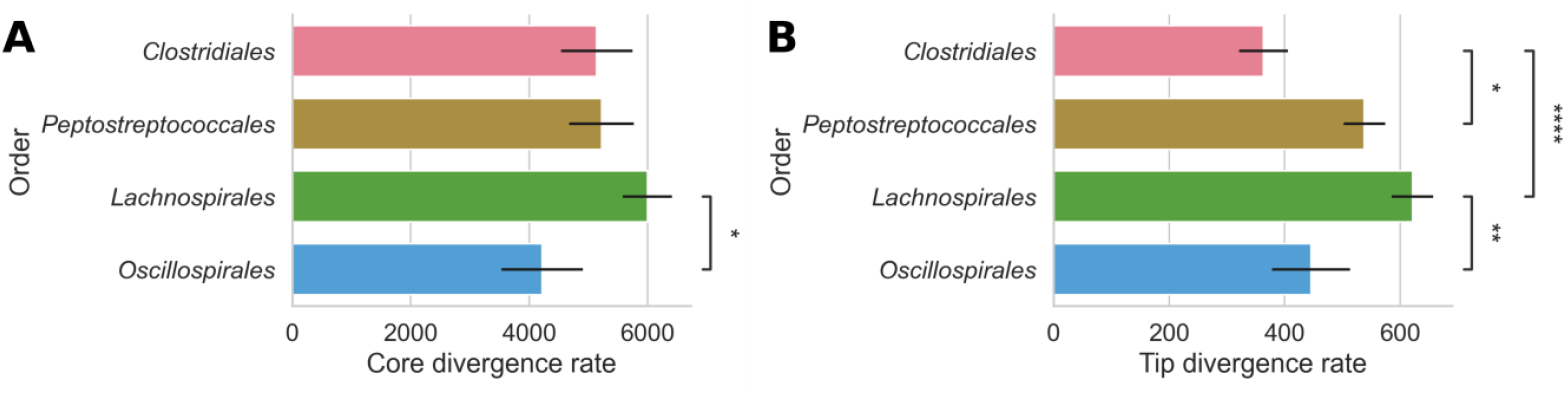
Fitted core (A) and tip (B) divergence rates of the GD models for each order, including standard error and statistical significance annotations. *Oscillospirales* have a significantly lower divergence rate, consistent with their host-associated nature, whereas *Lachnospirales* display a high divergence rate inconsistent with their host-associated status. This inconsistency can be explained by *Lachnospirales* tending to be in the earlier stages of host adaptation

### Ancestral state reconstruction uncovers different ancestral lifestyles for the host-associated orders

An ancestral state reconstruction (ASR) infers the clade-specific genes that have been passed on from a common ancestor to its descendants. By functionally annotating these reconstructed ancestral genes, it is possible to infer the lifestyle of this common ancestor, tracing the evolutionary origin of lifestyle differences. The ASR of the clostridial phylogeny shows a relatively high number of gene gain and loss events in the vicinity of the root node of each order (**Figure 4**).

**Figure 4.**
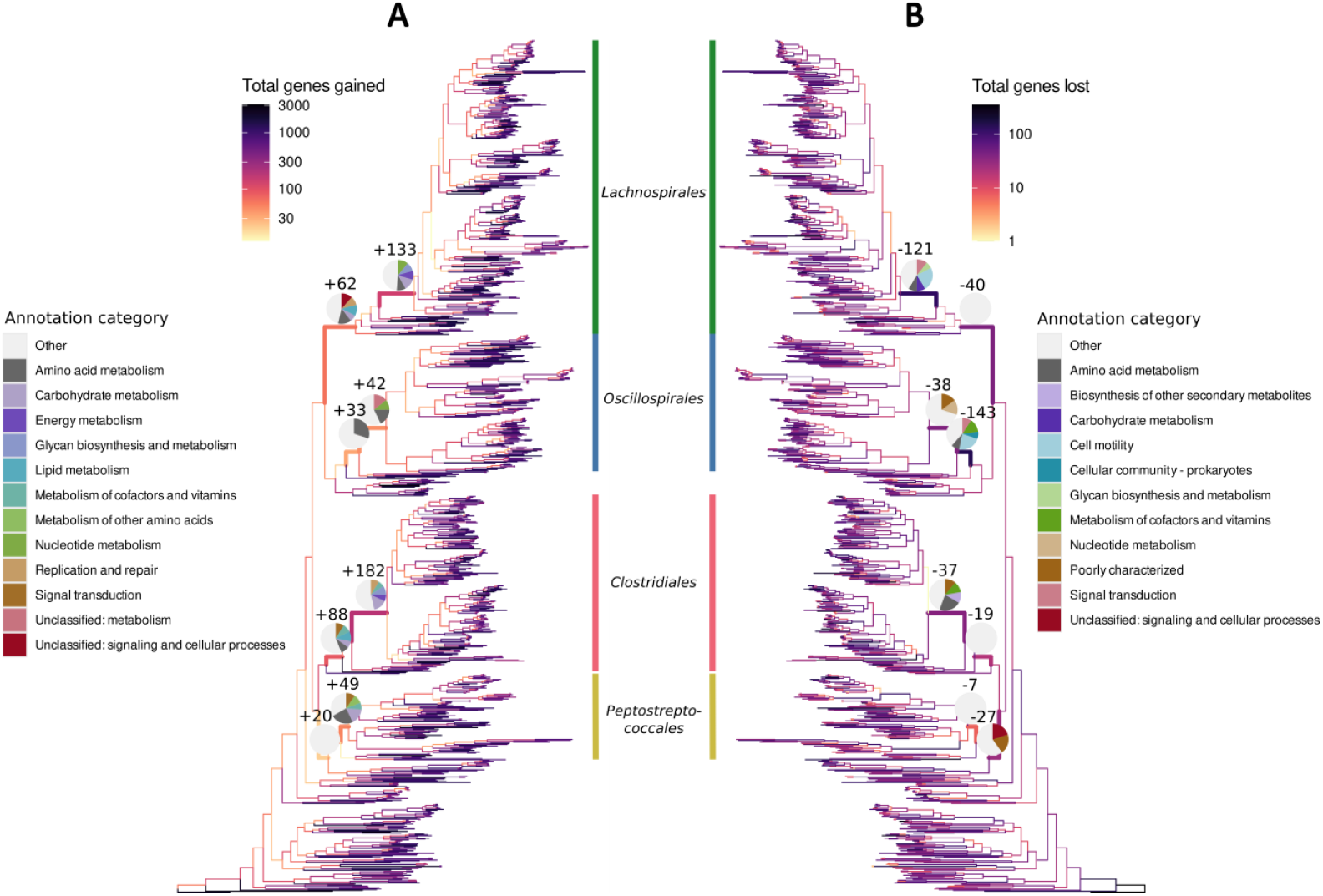
Wagner parsimony gene gain (A) and loss (B) profiles for the full genome set (coloured branches), and absolute number and KEGG BRITE level B annotation profile of exchanged genes for a selection of ancestral nodes (inset pie charts). For each node, only the top five categories are shown, each shown category requiring minimum three counts, and excluding the “Unannotated” category.

A total of 796 genes appear to have been present in the last common ancestor (LCA) of the *Clostridia*. These are associated with core cellular processes like glycolysis, tRNA biosynthesis and the ribosomes. In addition, they include genes for flagellar assembly, two-component signalling (2CS) systems and DNA repair systems, suggesting that the last common ancestor of the clostridial lineage was a well-equipped free-living microbe.

Analysis of the most recent common ancestors (MRCA) of each individual order interesting differences in both the number and functional annotation of the exchanged genes (**Add. File 5**). The MRCA of *Clostridiales* gained 88 genes, only half of which could be functionally annotated. These are mostly associated with metabolism, signal transduction and cell motility, consistent with a free-living lifestyle. In contrast, this MRCA lost only 19 genes, of which the annotated fraction is again mostly metabolism-related. Notably, the ancestor of a large subcluster containing the *Clostridium sensu stricto* clade has gained 182 genes, which are mostly associated with metabolism, and DNA replication and repair, indicating that this subcluster even further refined and expanded its ancestral free-living capacities.

The *Oscillospirales* MRCA gained 42 genes and lost 38, mostly related to metabolism in both cases. Interestingly, the direct ancestor of this node has lost a considerable number of gene families involved in cell motility, cofactor metabolism and signal transduction pathways. These losses align with the reduction in free-living capabilities that typically occurs when a microorganism transitions to a host-associated lifestyle.

In contrast, the *Lachnospirales* MRCA has gained 62 genes and lost 40. The acquired genes are associated with amino acid and lipid metabolism, and DNA replication and repair, while most lost ones lack functional annotation. Also here, a pronounced gene exchange was observed in a descendant lineage corresponding to the MRCA of the historical *Clostridium XIVa* clade, which gained 133 genes and lost 121. Gained functionalities affect several metabolic systems, including secondary metabolite biosynthesis, while lost functionalities are linked with cell motility and signal transduction. Both the number of exchanged genes and the combination of metabolic expansion with the loss of free-living traits suggest that the *Lachnospirales* are at a different stage of host association compared to the *Oscillospirales*.

Lastly, the *Peptostreptococcales* ancestral lineages exchanged relatively few genes. For example, the MRCA gained 49 genes and lost only 7. The gained genes are involved in several metabolic subsystems, such as amino acid metabolism and glycan biosynthesis. This low level of gene exchange may indicate that *Peptostreptococcales* are either at the end of the host conversion process or have only just started it.

### Focused mobilome analysis confirms different levels of host adaptation

Both the GD and the ASR analyses indicate varying levels of host association among the clostridial orders. Moreover, the higher metabolic versatility observed in *Lachnospirales* may be related to lower levels of host dependence. To further substantiate these inferred stages of host adaptation, we screened the entire genome dataset for the abundance of genomic hallmarks associated with progressing host adaptation.

Specifically, the transition from a free-living to a host-associated lifestyle in bacteria is frequently coupled with a genome streamlining process driven by insertion sequences (ISEs) and intrinsic mutations (Siguier, Gourbeyre, and Chandler 2014). ISes are a type of intracellular mobile genetic elements (MGEs) that employ transposases for their mobility. When bacteria move from a nutritionally poor free-living environment to a nutrient-rich host-associated niche, many genes become redundant due to the constant availability of metabolites. This causes a reduced selection pressure on these non-essential genes, enabling higher mutation rates and IS insertions, which, together, drive pseudogene formation (Marais, Calteau, and Tenaillon 2008). Over time, the accumulation and multiplication of IS regions promotes genome rearrangements and deletions, ultimately yielding smaller streamlined genomes tailored to new host-associated lifestyles (**Figure 5G**) (Siguier, Gourbeyre, and Chandler 2014).

**Figure 5.**
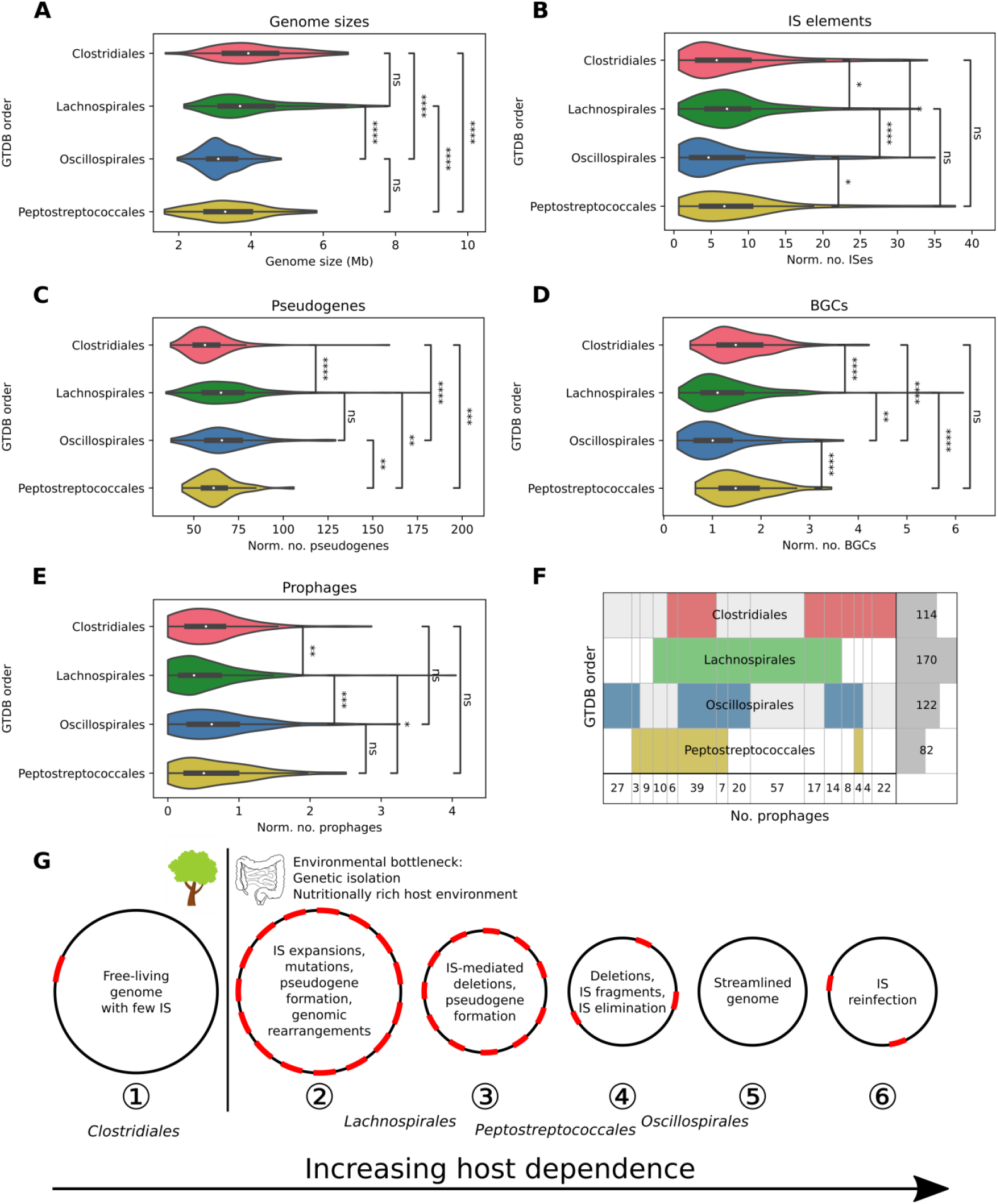
Screening of the genomic hallmarks of host dependence. **(A) – (F)** Comparison of the abundance of a genomic feature across all orders. Feature counts are normalised by dividing by the genome length in Mb. The applied statistical test is the Brunner-Munzel test (α = 0.05). (A) genome size, (B) insertion sequence elements, (C) pseudogenes, (D) biosynthetic gene clusters, (E) prophages, (F) Taxonomic order of the prophage hosts. **(G) Schematical overview of the impact of the genome streamlining process on the bacterial genomic architecture**, including the inferred position of each clostridial order in this process. Figure inspired on (Siguier, Gourbeyre, and Chandler 2014). (1) The (ancestral) free-living genome has only few ISes. (2) As a result of an environmental bottleneck, IS expansion occurs, leading to an increased abundance of ISes, pseudogenes and mutations. Genomic rearrangements occur. (3) With time, the repetitive regions of the ISes facilitate the deletion of non-essential genomic regions. (4) Eventually, extensive deletion leads to the elimination of the ISEs. (5) Finally, the genome reaches its streamlined form, with a gene content tailored for life in its host environment. (6) Upon contact with IS-carrying strains or phages, ISes may be reintroduced.

To test this hypothesis, we screened the complete genome dataset for the abundance of multiple MGEs and genomic features associated with host adaptation: (1) genome sizes, (2) ISes, (3) pseudogenes, (4) prophage regions, and (5) biosynthetic gene clusters. (1), (2) and (3) are established hallmarks of the genome streamlining process. (4) may reflect a shift in environmental niche. (5) may provide an indication of metabolic versatility, as it depends on the availability of diverse precursor metabolites.

The results align strikingly well with our previous interpretations. First, *Oscillospirales* and *Peptostreptococcales* genomes are significantly smaller than those of *Clostridiales* and *Lachnospirales* (**Figure 5A**).

Second, *Lachnospirales* and *Peptostreptococcales* genomes harbour significantly more ISes than those of *Clostridiales*, which, in turn, contain more than *Oscillospirales* genomes (**Figure 5B**). This pattern is consistent with the hypothesis that *Lachnospirales* are in the early stages of host adaptation and are thus undergoing IS expansion. Furthermore, the smaller genomes and lower number of ISes in *Peptostreptococcales* suggest that this latter order is at a slightly more advanced stage of the host transition process than the *Lachnospirales*.

A more detailed analysis of the IS families revealed some substantial differences in their distribution across the clostridial orders (**Add. File 2: Figure S6**). For example, family IS1634 is only present in *Lachnospirales* and *Peptostreptococcales*, while family IS5 is only found in *Oscillospirales*. Notably, families IS110, IS66 and ISL3 are more prevalent in *Lachnospirales* species. As these IS families do not have a specific insertion site preference (Siguier, Gourbeyre, and Chandler 2014; Siguier et al. 2015), their abundances may merely be the result of IS expansion during the host conversion process.

Third, pseudogenes are significantly less abundant in *Clostridiales* than in *Peptostreptococcales*, which in turn harbour less than *Oscillospirales* and *Lachnospirales*. (**Figure 5C**). This implies that both gut-associated clusters have undergone or are undergoing more intense pseudogene formation, consistent with progressing genome streamlining during host adaptation.

Fourth, the number of secondary metabolite BGCs is significantly higher in *Clostridiales* and *Peptostreptococcales*, with *Lachnospirales* and *Oscillospirales* at comparably lower levels (**Figure 5D**). This suggests that *Clostridiales* and *Peptostreptococcales* species maintain a greater capacity for specialised metabolism, whereas the other two orders appear to have lost a considerable fraction of their biosynthetic capacities. Given that other genomic features indicate that *Lachnospirales* are at an early stage of host adaptation, this observation may imply that BGCs are among the first genomic features to be eliminated, even when metabolic versatility is maintained. Alternatively, this may also reflect current limitations in the detection and annotation of BGCs from gut-associated species. Analysis of BGC classes revealed that the most substantial differences in abundance occur among the cyclic lactone autoinducer (CLA), NRPS(-like), and RiPP-like classes. In addition, there is a consistent presence of ranthipeptides and terpene precursors across all orders (**Add. File 2: Figure S7**).

Finally, *Lachnospirales* genomes contain significantly less prophages than those of the other orders (**Figure 5E**). In terms of host range, however, *Lachnospirales* carried the highest diversity in prophage types, while *Peptostreptococcales* the lowest. Interestingly, about 75% of the detected prophage types were found in clostridial species that were part of different orders, with 39 occurring across all four orders (**Figure 5F**). In addition, some prophages carry passenger genes for methyltransferases, restriction enzymes, toxin(-antitoxin system)s and transporters (including beta-lactam efflux pumps and other resistance proteins), highlighting a mechanism by which clostridial species may acquire new genes and adaptive traits (**Add. File 6**). Although clear differences in prophage abundance and diversity were observed, it remains unclear whether this merely reflects a niche shift or whether these prophage-encoded genes that have become essential for the bacterium’s survival due to functional integration.

In conclusion, the abundance patterns of these genomic features further support the notion that the clostridial species orders are in various stages of the host adaptation process (**Figure 5G**).

### BGC clustering analysis reveals strongly conserved BGCs

The species in our clostridial dataset have faced diverse ecological and evolutionary selection pressures. Hence, BGCs that have remained conserved across these varying conditions, likely encode functions that offer substantial evolutionary advantages that justify their long-term conservation. Such high-potential BGCs are expected to emerge in BGC clustering analyses.

Our clustering analysis revealed that the five largest BGC groups consist of two terpene-precursor groups and three ranthipeptide groups. Both *Clostridiales* and *Lachnospirales* harbour one terpene-precursor and one or two ranthipeptide BGC groups (**Figure 6**). Alignment of representative gene clusters from the three largest ranthipeptide groups shows that the core biosynthetic genes, i.e. the SCIFF precursor and the RRE gene, are conserved across all groups (**Add. File 2: Figure S8)**. Moreover, these ranthipeptide BGCs are well-distributed across the phylogeny, with each species encoding maximum one ranthipeptide (**Add. File 2: Figure S9**). Interestingly, two of these ranthipeptide BGCs have been experimentally shown to participate in quorum sensing activities (Chen et al. 2020). The fact that these two BGCs belong to the same ranthipeptide group in our analysis suggests that similar quorum sensing functions may be associated with this entire ranthipeptide group.

**Figure 6.**
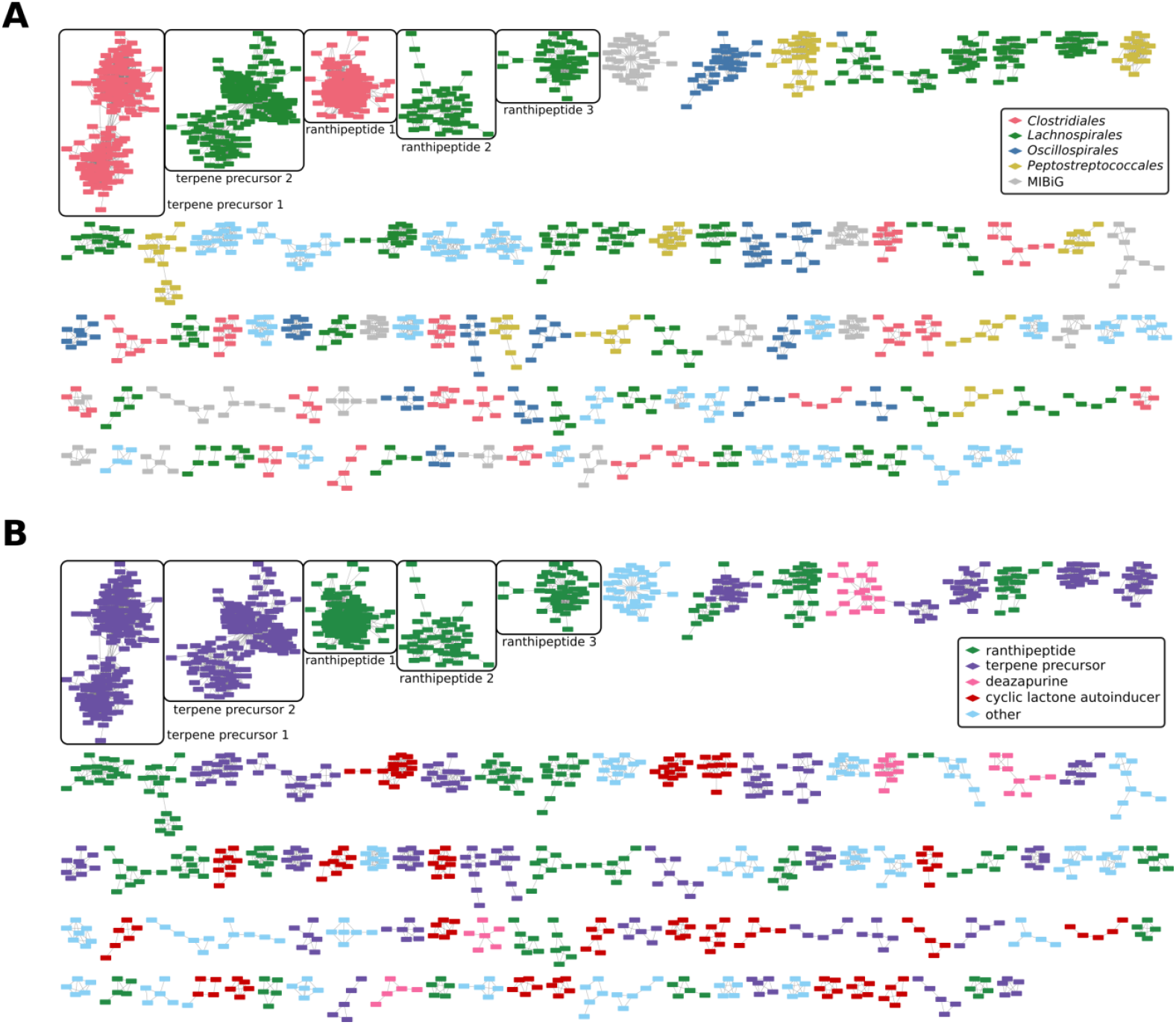
BiG-SCAPE similarity networks of all BGCs detected in our full genome set (threshold 0.3), annotated (A) by clostridial order or as MIBiG entry, and (B) by BGC type. The five largest groups on the top left are either terpene precursors or ranthipeptides. Only groups with at least five members are shown.

The smaller identified BGC groups are precited to encode cyclic lactone autoinducers and deazapurines, next to more smaller ranthipeptides and terpene precursor groups. We co-clustered our BGC set with the MIBiG database, a highly curated and standardised repository of known BGCs (Zdouc et al. 2025). Interestingly, very few MIBiG BGCs co-cluster with our clostridial BGCs, underscoring the novelty of these conserved ranthipeptide and terpene precursor BGCs, and their high potential for functional significance. We also observed a large group of singleton BGCs (**Add. File 7**), which may comprise BGCs acquired more recently by horizontal gene transfer or gene rearrangement. While these singletons have not undergone sufficient evolutionary selection to refine their functional significance, the possibility that they encode potent and biologically relevant activities cannot be excluded.

## Discussion

This work presents the first comparative pangenomics analysis of multiple orders of the *Clostridia* class and reveals intriguing differences in their host adaptation tendencies. Moreover, it uncovered a remarkably large repertoire of uncharacterised BGCs, including several highly conserved ranthipeptide families, highlighting substantial untapped biosynthetic potential.

A major finding of this study is that the clostridial orders are at varying stages of host adaptation. By using multiple independent genomics approaches in parallel, we were able to position each order on the host adaptation spectrum. In particular, the early-stage host association trend of the *Lachnospirales* adds an important nuance to their commonly assigned status as host-associated gut symbionts. Much of the genomic and metabolic diversity of this clade is obscured by the generic label “host-associated”. Bacteria in the early stages of host association likely still retain part of their ancestral free-living gene repertoire, a hypothesis supported by our KEGG pathway completeness assessment, which revealed higher metabolic versatility among *Lachnospirales* species. The combination of their higher innate capacity to produce more complex metabolites and their adaptation towards host environments positions these species as attractive starting points for the development of anaerobic chassis organisms for heterologous expression of complex biosynthetic pathways. Moreover, their ecological compatibility with the gut environment suggests a high potential for therapeutic applications, such as delivery of bioactive compounds or engineered functions in gut microbiomes (Kubiak and Minton 2015).

Another key outcome of this study is the comprehensive identification and comparative analysis of clostridial BGCs, providing, to the best of our knowledge, the first comparative view of the natural product repertoire beyond PKS and NRPS classes across multiple clostridial orders. Despite extensive host adaptation and a long track record of exposure to varying selection pressures, several ranthipeptide and terpene precursor BGCs have been widely conserved. The lack of MIBiG entries co-clustering with these BGCs underscores their novelty, tapping further into the unexplored richness of BGCs from the anaerobic realm. These novel conserved BGCs are prime targets for further experimental characterisation.

Nevertheless, several considerations should be taken into account when interpreting these findings. First, as the study conducted a high-level analysis averaging counts from taxonomic orders, the observed trends reflect general evolutionary tendencies of each order and should not be extrapolated to individual species without further analysis. For example, it may be possible that a *Lachnospirales* species is already highly adapted to its host because of higher selection pressures, or because it started the host adaptation process early compared to its close relatives. Second, we dereplicated our dataset to ensure an equal species representation in the pangenome. Non-representative genomes may harbour even more novel diversity, both at the pangenome level as well as at the BGC level.

Overall, the combined application of comparative genomics, phylogenetic techniques and targeted analysis of genomic features at higher taxonomic levels offers a flexible approach to investigate host adaptation trends. We uncovered counterintuitive heterogeneity in host adaptation among clostridial species clusters, which translates into evolving genome architectures and different levels of metabolic versatility. Taken together, the findings of this study are an encouraging starting point for the systematic investigation of adaptation levels in other “host-associated” bacteria classes than *Clostridia*. Moreover, in the light of natural product research, our findings pave the way for (1) establishing clostridial species as new anaerobic host organisms for a diverse range of applications, as well as for (2) expanding the natural product repertoire further into the anaerobic realm.

## Supporting information

additional_file1_genomes_metadata

additional_file3_pangenome_parameter_sensitivity

additional_file4_BiG-SCAPE_threshold_sensitivity

additional_file5_ASR_annotation

additional_file6_passenger_genes_prophages

additional_file7_BiG-SCAPE_cytoscape_sessions

additional_file8_COG_fractions_and_counts

## Data availability

All genome assemblies analysed in this study were retrieved from NCBI RefSeq. Accession codes and relevant metadata can be found in Add. File 1.

## Code availability

The full analysis pipeline has been deposited at GitHub (https://github.com/LucoDevro/closHomer) and at Zenodo (https://doi.org/10.5281/zenodo.14719962).

## Author contribution statement

LDV and JM conceived this study. LDV and JM designed the research. LDV performed the data analysis, implemented the workflow, and gathered the results. LDV analysed and interpreted the data, with assistance from JA and JM. LDV wrote the article, which was revised by LDV, JA and JM.

## Funding statement

This work was funded by the European Commission through the European Research Council (MiStiC, 101078461).

## Competing interest statement

We declare no competing interests.

## Notes

https://github.com/LucoDevro/closHomer

https://doi.org/10.5281/zenodo.14719962

### Competing Interest Statement

The authors have declared no competing interest.

### Summary of Updates

Final tweaks (language polishing, tuning down novelty statement, adding new supplementary file and referring to it in caption of Figure 1, clarifying selected categories in Figure 4)

## References

Asnicar, Francesco, et al. (2020), ‘Precise Phylogenetic Analysis of Microbial Isolates and Genomes from Metagenomes Using PhyloPhlAn 3.0’, Nature Communications, 11/1: 2500, 10.1038/s41467-020-16366-7.

Behnken, Swantje, and Christian Hertweck (2012), ‘Anaerobic Bacteria as Producers of Antibiotics’, Applied Microbiology and Biotechnology, 96/1: 61–7, 10.1007/s00253-012-4285-8.

Birgen, Cansu, et al. (2021), ‘Butanol Production from Lignocellulosic Sugars by Clostridium Beijerinckii in Microbioreactors’, Biotechnology for Biofuels, 14/1, 10.1186/s13068-021-01886-1.

Blin, Kai, et al. (2025), ‘AntiSMASH 8.0: Extended Gene Cluster Detection Capabilities and Analyses of Chemistry, Enzymology, and Regulation’, Nucleic Acids Research (published online 25 April), 10.1093/nar/gkaf334.

Buchfink, Benjamin, Klaus Reuter, and Hajk-Georg Drost (2021), ‘Sensitive Protein Alignments at Tree-of-Life Scale Using DIAMOND’, Nature Methods, 18/4: 366–8, 10.1038/s41592-021-01101-x.

Cantalapiedra, Carlos P, et al. (2021), ‘EggNOG-Mapper v2: Functional Annotation, Orthology Assignments, and Domain Prediction at the Metagenomic Scale’, Molecular Biology and Evolution, 38/12: 5825–9, 10.1093/molbev/msab293.

Capella-Gutiérrez, Salvador, José M. Silla-Martínez, and Toni Gabaldón (2009), ‘TrimAl: A Tool for Automated Alignment Trimming in Large-Scale Phylogenetic Analyses’, Bioinformatics, 25/15: 1972–3, 10.1093/bioinformatics/btp348.

Chaumeil, Pierre-Alain, et al. (2022), ‘GTDB-Tk v2: Memory Friendly Classification with the Genome Taxonomy Database’, Bioinformatics, 38/23: 5315–6, 10.1093/bioinformatics/btac672.

Chen, Yunliang, et al. (2020), ‘The SCIFF-Derived Ranthipeptides Participate in Quorum Sensing in Solventogenic Clostridia’, Biotechnology Journal, 15/10, 10.1002/biot.202000136.

Cohen, Jonathan E., et al. (2017), ‘Comparative Pathogenomics of Clostridium Tetani’, PLOS ONE, 12/8: e0182909, 10.1371/journal.pone.0182909.

Cruz-Morales, Pablo, et al. (2019), ‘Revisiting the Evolution and Taxonomy of Clostridia, a Phylogenomic Update’, Genome Biology and Evolution, 11/7: 2035–44, 10.1093/gbe/evz096.

Csurös, Miklós (2010), ‘Count: Evolutionary Analysis of Phylogenetic Profiles with Parsimony and Likelihood’, Bioinformatics, 26/15: 1910–2, 10.1093/bioinformatics/btq315.

Dufresne, Alexis, Laurence Garczarek, and Frédéric Partensky (2005), ‘Accelerated Evolution Associated with Genome Reduction in a Free-Living Prokaryote’, Genome Biology, 6/2: R14, 10.1186/gb-2005-6-2-r14.

Gautreau, Guillaume, et al. (2020), ‘PPanGGOLiN: Depicting Microbial Diversity via a Partitioned Pangenome Graph’, PLOS Computational Biology, 16/3: e1007732, 10.1371/journal.pcbi.1007732.

Gilchrist, Cameron L.M., and Yit Heng Chooi (2021), ‘Clinker & Clustermap.Js: Automatic Generation of Gene Cluster Comparison Figures’, Bioinformatics, 37/16: 2473–5, 10.1093/bioinformatics/btab007.

Katoh, K., and D. M. Standley (2013), ‘MAFFT Multiple Sequence Alignment Software Version 7: Improvements in Performance and Usability’, Molecular Biology and Evolution, 30/4: 772–80, 10.1093/molbev/mst010.

Kubiak, Aleksandra M., and Nigel P. Minton (2015), ‘The Potential of Clostridial Spores as Therapeutic Delivery Vehicles in Tumour Therapy’, Research in Microbiology, 166/4: 244–54, 10.1016/j.resmic.2014.12.006.

Letunic, Ivica, and Peer Bork (2024), ‘Interactive Tree of Life (ITOL) v6: Recent Updates to the Phylogenetic Tree Display and Annotation Tool’, Nucleic Acids Research, 52/W1: W78–82, 10.1093/nar/gkae268.

Letzel, Anne Catrin, Sacha J. Pidot, and Christian Hertweck (2013), ‘A Genomic Approach to the Cryptic Secondary Metabolome of the Anaerobic World’, Natural Product Reports, 30/3: 392–428, 10.1039/c2np20103h.

Marais, Gabriel A.B., Alexandra Calteau, and Olivier Tenaillon (2008), ‘Mutation Rate and Genome Reduction in Endosymbiotic and Free-Living Bacteria’, Genetica, 134/2: 205–10, 10.1007/s10709-007-9226-6.

Maturana, José Luis, and Juan P. Cárdenas (2021), ‘Insights on the Evolutionary Genomics of the Blautia Genus: Potential New Species and Genetic Content Among Lineages’, Frontiers in Microbiology, 12, 10.3389/fmicb.2021.660920.

Minh, B. Q., M. A. T. Nguyen, and A. von Haeseler (2013), ‘Ultrafast Approximation for Phylogenetic Bootstrap’, Molecular Biology and Evolution, 30/5: 1188–95, 10.1093/molbev/mst024.

Minh, Bui Quang, et al. (2020), ‘IQ-TREE 2: New Models and Efficient Methods for Phylogenetic Inference in the Genomic Era’, Molecular Biology and Evolution, 37/5: 1530–4, 10.1093/molbev/msaa015.

Navarro-Muñoz, Jorge C., et al. (2020), ‘A Computational Framework to Explore Large-Scale Biosynthetic Diversity’, Nature Chemical Biology, 16/1: 60–8, 10.1038/s41589-019-0400-9.

O’Leary, Nuala A., et al. (2024), ‘Exploring and Retrieving Sequence and Metadata for Species across the Tree of Life with NCBI Datasets’, Scientific Data, 11/1: 732, 10.1038/s41597-024-03571-y.

Pahalagedara, Amila Srilal Nawarathna Weligala, et al. (2020), ‘Antimicrobial Production by Strictly Anaerobic Clostridium Spp.’, International Journal of Antimicrobial Agents, 55/5, 10.1016/j.ijantimicag.2020.105910.

Pascal Andreu, Victòria, et al. (2023), ‘GutSMASH Predicts Specialized Primary Metabolic Pathways from the Human Gut Microbiota’, Nature Biotechnology, 41/10: 1416–23, 10.1038/s41587-023-01675-1.

Patakova, Petra, et al. (2019), ‘Acidogenesis, Solventogenesis, Metabolic Stress Response and Life Cycle Changes in Clostridium Beijerinckii NRRL B-598 at the Transcriptomic Level’, Scientific Reports, 9/1, 10.1038/s41598-018-37679-0.

Roberts, Seth B, et al. (2010), ‘Genome-Scale Metabolic Analysis of Clostridium Thermocellum for Bioethanol Production’, BMC Systems Biology, 4/1: 31, 10.1186/1752-0509-4-31.

Ruiz-Perez, Carlos A., Roth E. Conrad, and Konstantinos T. Konstantinidis (2021), ‘MicrobeAnnotator: A User-Friendly, Comprehensive Functional Annotation Pipeline for Microbial Genomes’, BMC Bioinformatics, 22/1: 11, 10.1186/s12859-020-03940-5.

Salamzade, Rauf, Aamuktha Kottapalli, and Lindsay R. Kalan (2025), ‘SkDER and CiDDER: Two Scalable Approaches for Microbial Genome Dereplication’, Microbial Genomics, 11/7, 10.1099/mgen.0.001438.

Sandhu, Brindar K., and Shonna M. McBride (2018), ‘Clostridioides Difficile’, Trends in Microbiology, 26/12: 1049–50, 10.1016/j.tim.2018.09.004.

Schwengers, Oliver, et al. (2021), ‘Bakta: Rapid and Standardized Annotation of Bacterial Genomes via Alignment-Free Sequence Identification’, Microbial Genomics, 7/11, 10.1099/mgen.0.000685.

Siguier, Patricia, et al. (2015), ‘Everyman’s Guide to Bacterial Insertion Sequences’, Microbiology Spectrum, 3/2, 10.1128/microbiolspec.mdna3-0030-2014.

Siguier, Patricia, Edith Gourbeyre, and Mick Chandler (2014), ‘Bacterial Insertion Sequences: Their Genomic Impact and Diversity’, FEMS Microbiology Reviews, 38/5: 865–91, 10.1111/1574-6976.12067.

Smith, Theresa J., Karen K. Hill, and Brian H. Raphael (2015), ‘Historical and Current Perspectives on Clostridium Botulinum Diversity’, Research in Microbiology, 166/4: 290–302, 10.1016/j.resmic.2014.09.007.

Sugimoto, Yuki, et al. (2019), ‘A Metagenomic Strategy for Harnessing the Chemical Repertoire of the Human Microbiome’, Science, 366/6471, 10.1126/science.aax9176.

Syberg-Olsen, Mitchell J, et al. (2022), ‘Pseudofinder: Detection of Pseudogenes in Prokaryotic Genomes’, Molecular Biology and Evolution, 39/7, 10.1093/molbev/msac153.

Tonkin-Hill, Gerry, et al. (2023), ‘Robust Analysis of Prokaryotic Pangenome Gene Gain and Loss Rates with Panstripe’, Genome Research, 33/1: 129–40, 10.1101/gr.277340.122.

Udaondo, Zulema, Estrella Duque, and Juan Luis Ramos (2017), ‘The Pangenome of the Genus Clostridium’, Environmental Microbiology, 19/7: 2588–603, 10.1111/1462-2920.13732.

Vieira-Silva, Sara, et al. (2016), ‘Species-Function Relationships Shape Ecological Properties of the Human Gut Microbiome’, Nature Microbiology, 1/8, 10.1038/nmicrobiol.2016.88.

Wishart, David S, et al. (2023), ‘PHASTEST: Faster than PHASTER, Better than PHAST’, Nucleic Acids Research, 51/W1: W443–50, 10.1093/nar/gkad382.

Xie, Zhiqun, and Haixu Tang (2017), ‘ISEScan: Automated Identification of Insertion Sequence Elements in Prokaryotic Genomes’, Bioinformatics, 33/21: 3340–7, 10.1093/bioinformatics/btx433.

Yu, Guangchuang, et al. (2017), ‘Ggtree: An R Package for Visualization and Annotation of Phylogenetic Trees with Their Covariates and Other Associated Data’, Methods in Ecology and Evolution, 8/1: 28–36, 10.1111/2041-210X.12628.

Zdouc, Mitja M, et al. (2025), ‘MIBiG 4.0: Advancing Biosynthetic Gene Cluster Curation through Global Collaboration’, Nucleic Acids Research, 53/D1: D678–90, 10.1093/nar/gkae1115.

