## additional_file2_supplementary_figures for "Comparative pangenomics unveils distinct host adaptation levels and conserved biosynthetic potential in microbiome *Clostridia*"

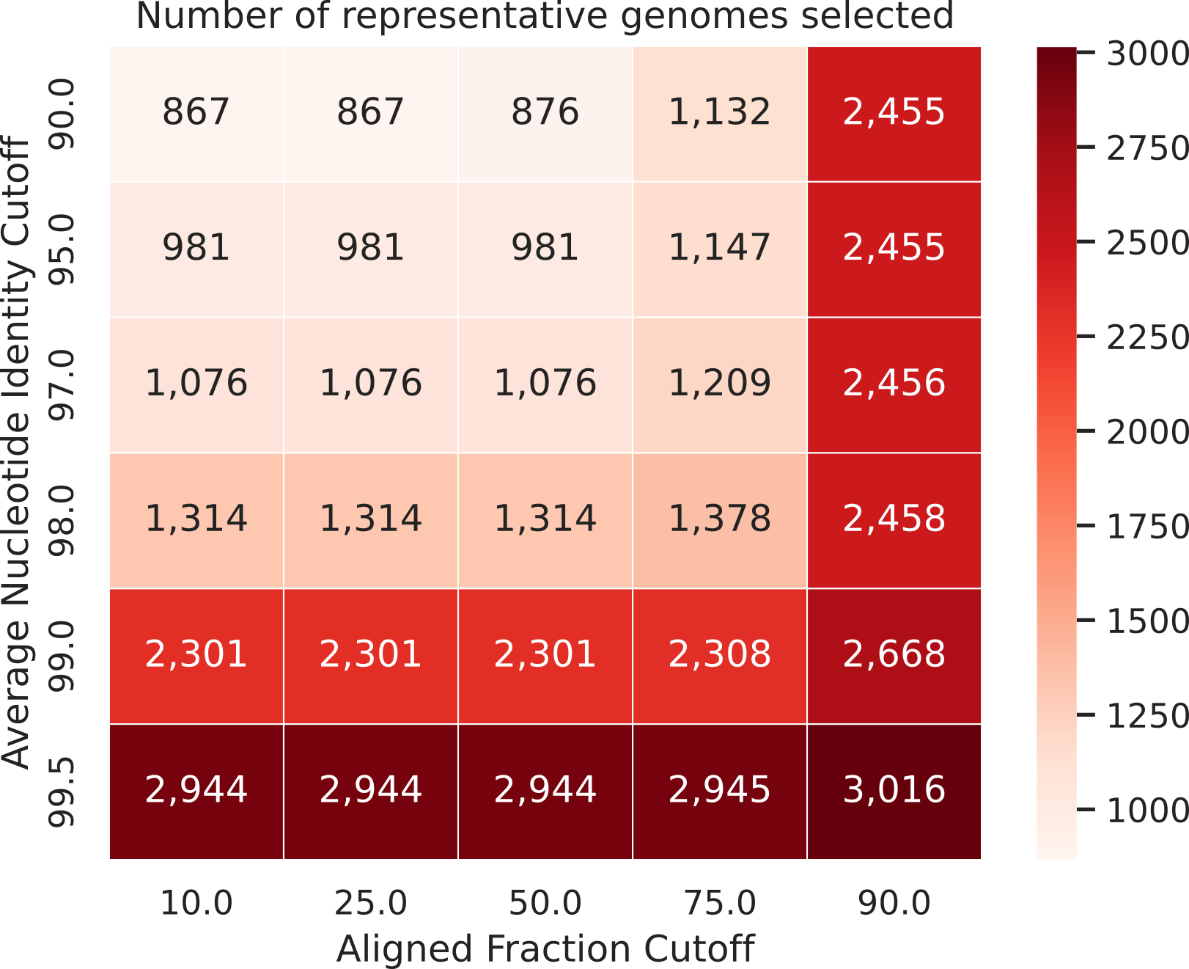

**Figure S1.** Parameter sensitivity analysis for the genome dereplication using skDER. The sharp increases in representative genomes suggest selecting an ANI cutoff around 97%, and an AF cutoff of 50%. Eventually, we selected an AF cutoff of 50%, and an ANI cutoff of 96%, consistent with the ANI threshold commonly used for species delineation.

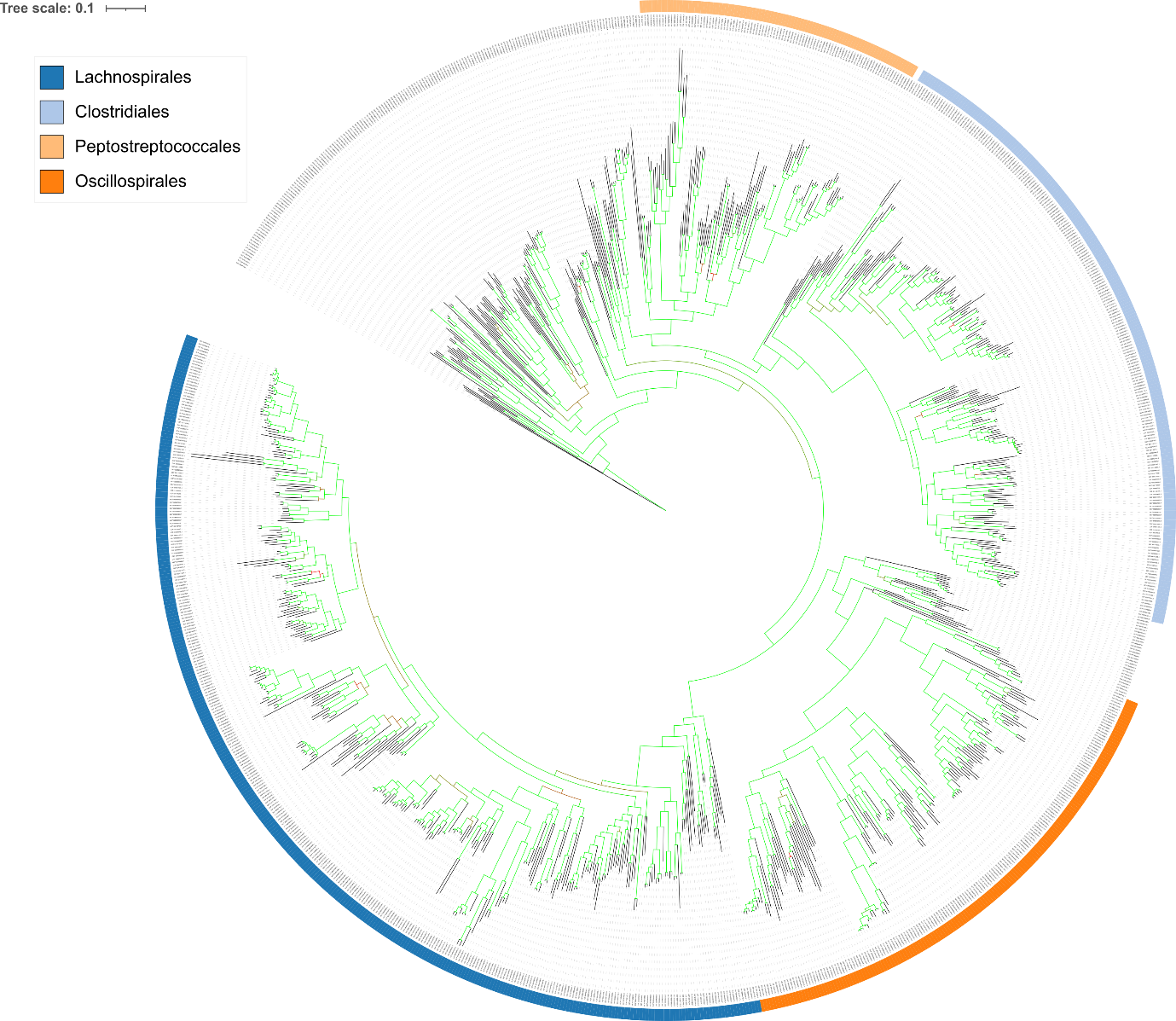

**Figure S2.** Rooted phylogeny annotated by taxonomic order according to GTDB (colour ring). Together, the four selected clostridial orders cover a large part of the Clostridia class.

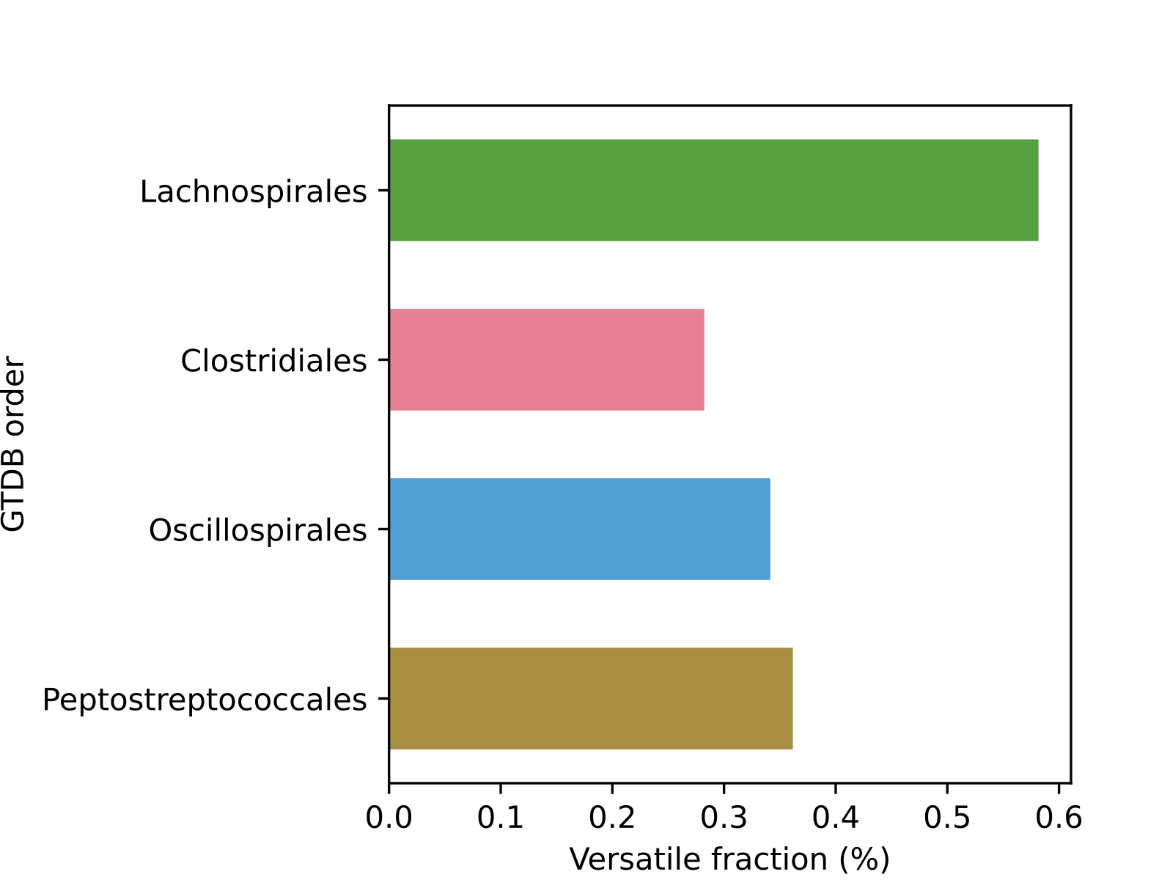

**Figure S3.** Fraction of the species in each order that is metabolically versatile according to the hierarchical clustering of the KEGG completeness analysis. The Lachnospirales have a remarkable high fraction compared to the other orders.

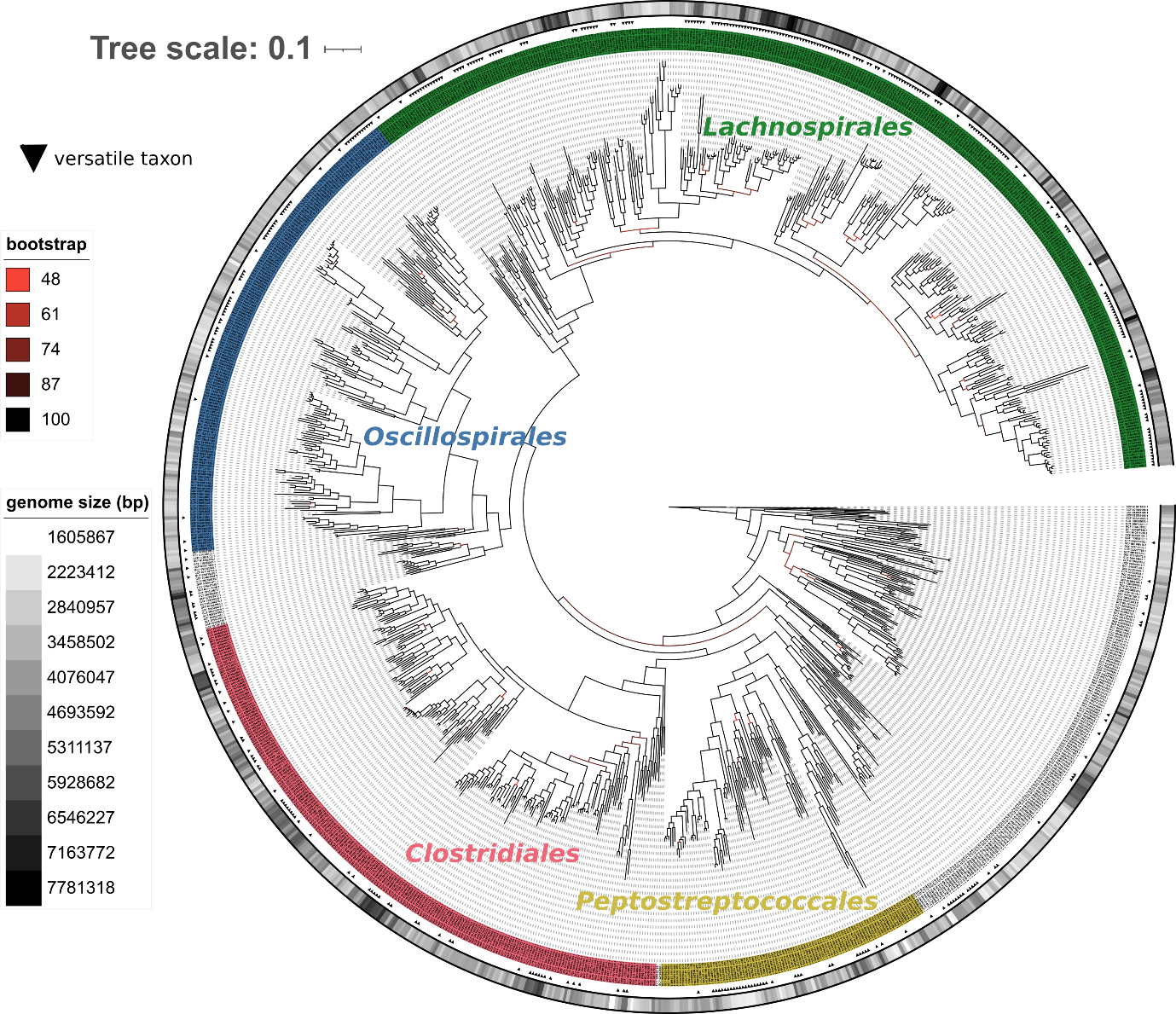

**Figure S4.** Rooted phylogeny annotated by taxonomic order (label colour), bootstrap support (branch colour), genome size (colour gradient ring) and high metabolic versatility (black triangular markers). Metabolically versatility does not cluster well with phylogenetic descent nor with genome size (except for Clostridiales), yet some lineages are consistently versatile.

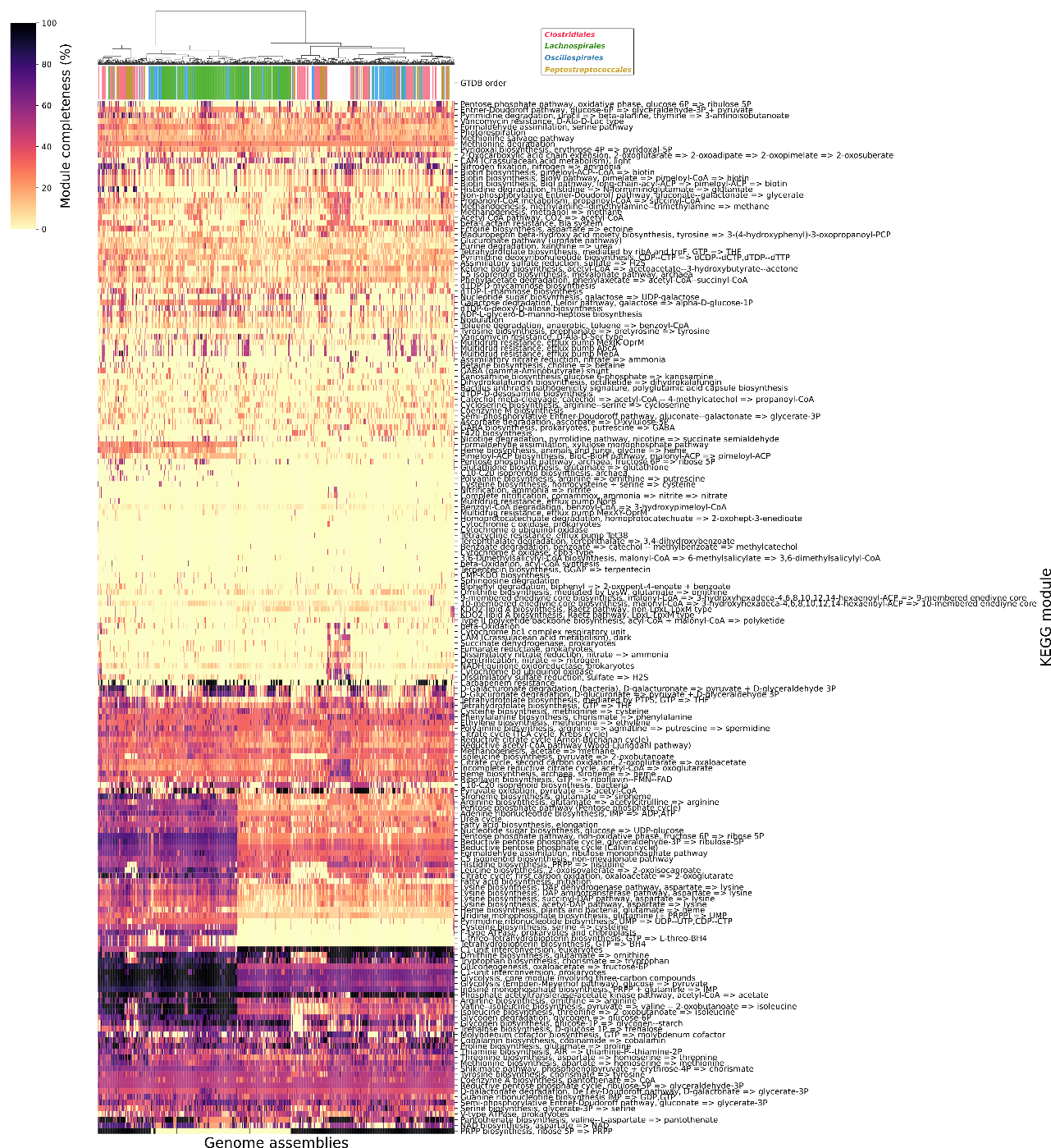

**Figure S5.** Ward’s clustered heatmap of KEGG module completeness, annotated by taxonomic order (top) and KEGG module (right).

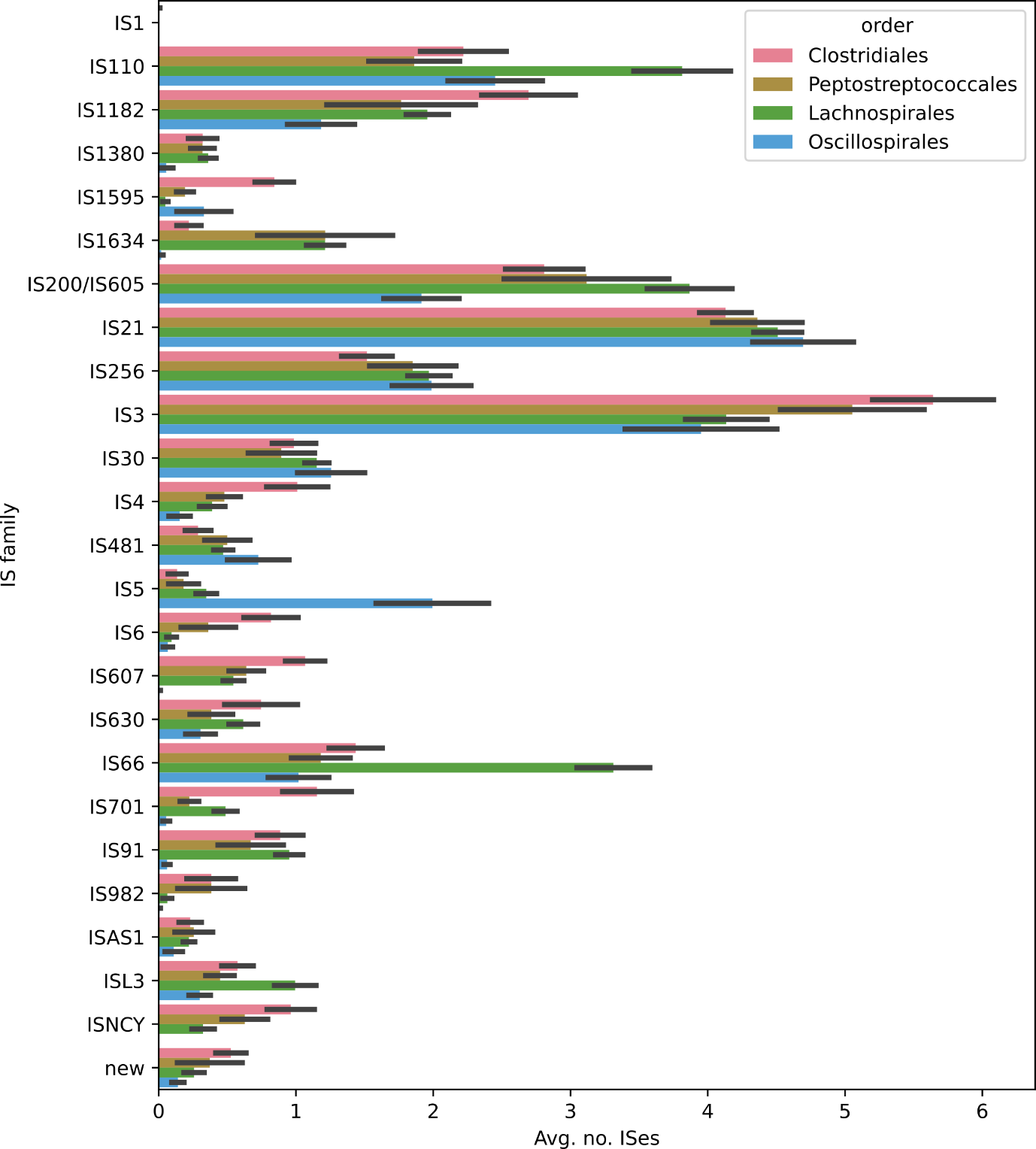

**Figure S6**. Average number of IS family members present in each taxonomic order, including standard error of the mean.

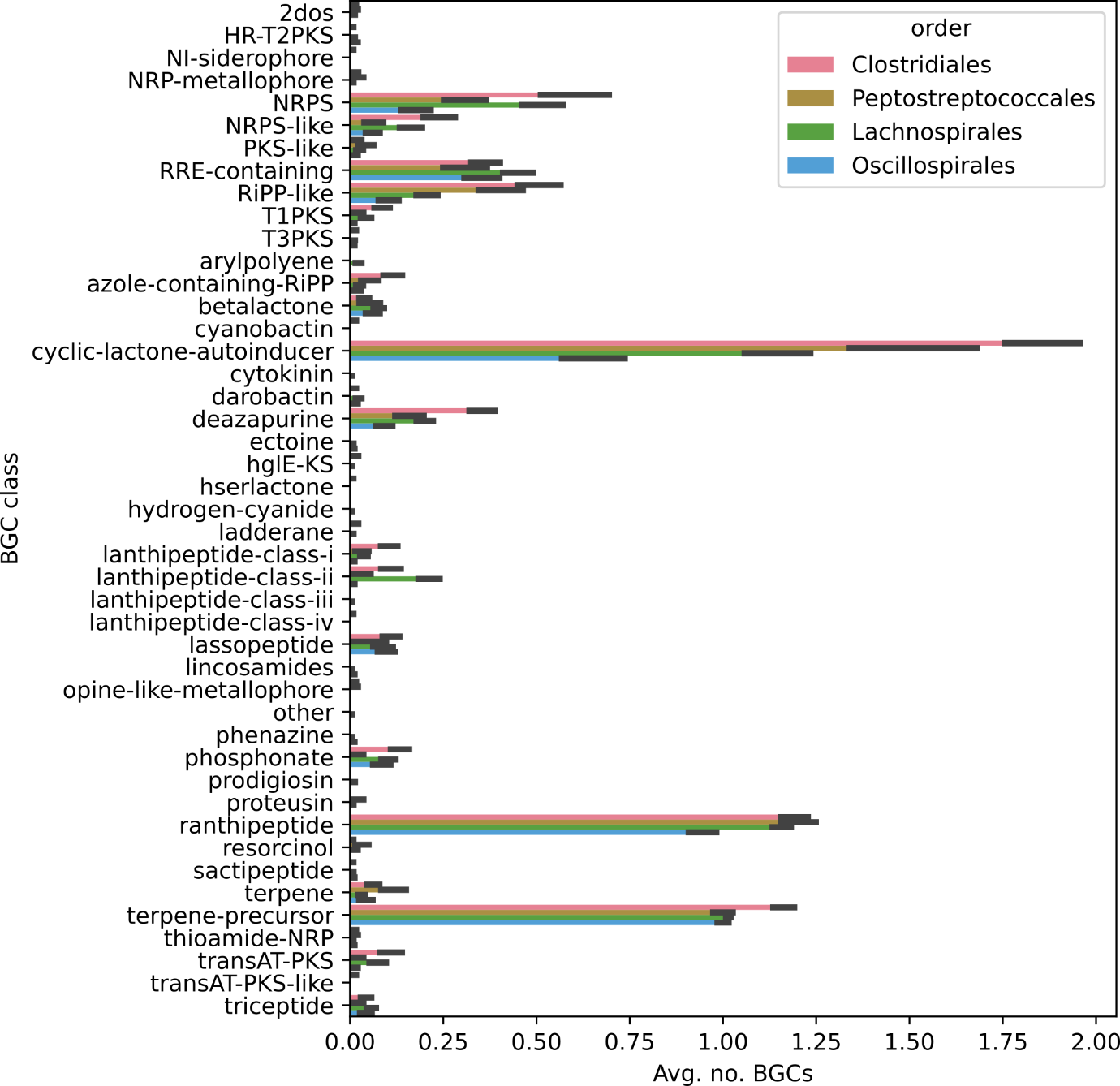

**Figure S7.** Average number of BGC types present in each taxonomic order, including standard error of the mean.

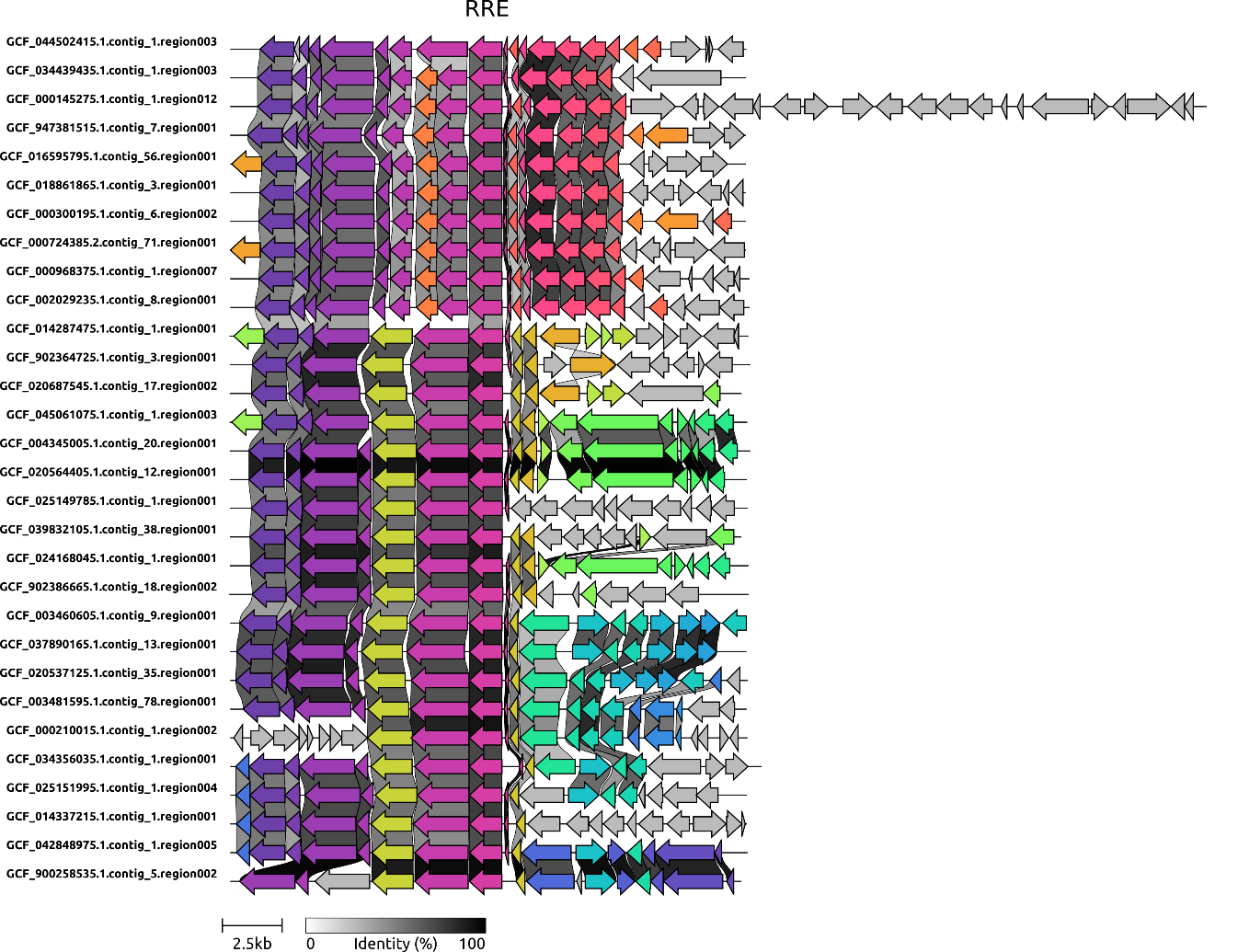

**Figure S8**. BGC gene alignment of a selection of ranthipeptide BGCs from the three biggest ranthipeptide BGC groups. The RRE gene (and the SCIFF precursor to the right, if detected) are well conserved across all three groups.

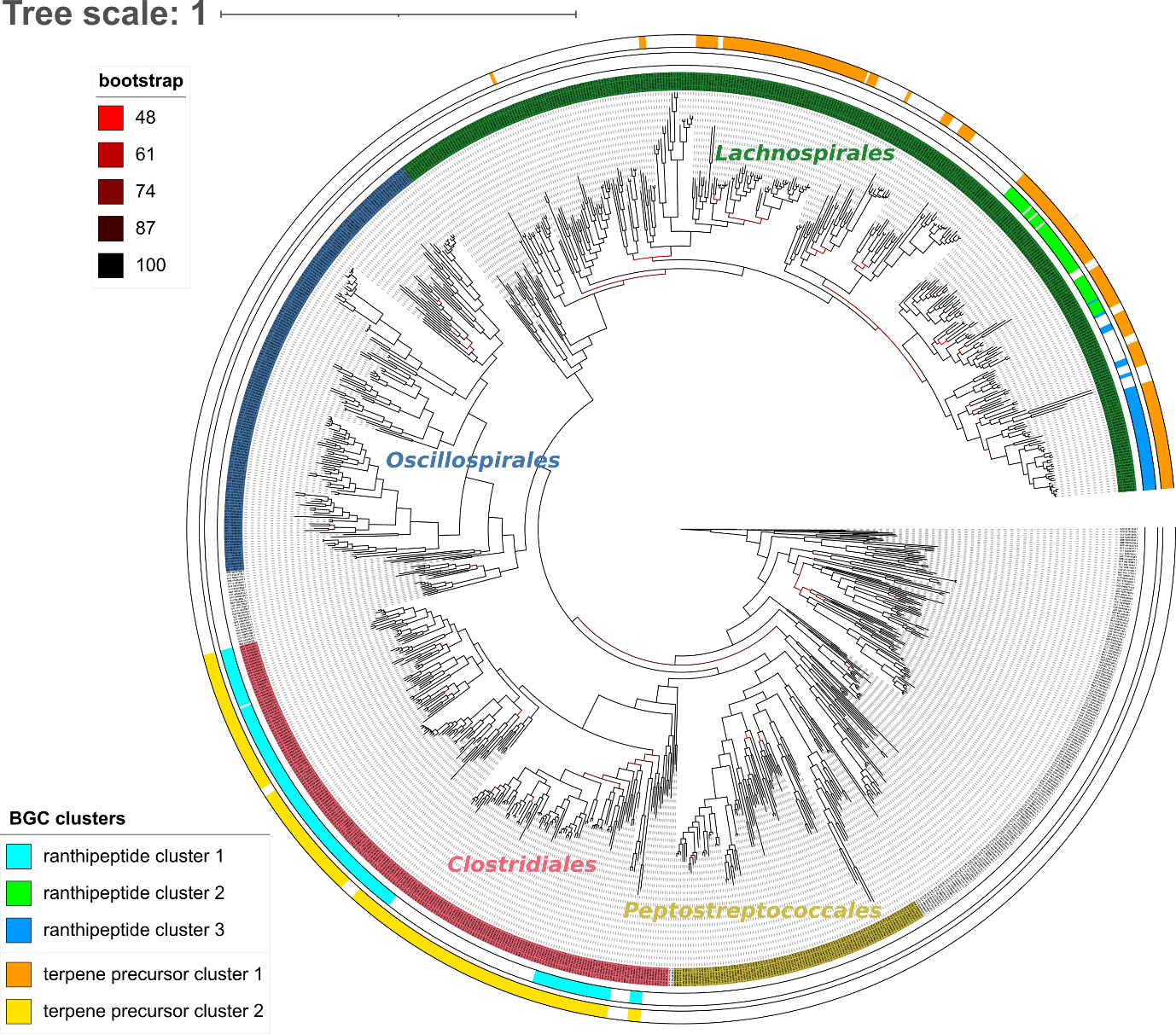

**Figure S9.** Rooted core genome phylogeny of the full clostridial genome set annotated by taxonomic order and by BGC group for the five biggest BGC group (three ranthipeptide and two terpene precursor groups).
